# Crystallographic fragment screening of CDK2-cyclin A: FragLites map sites of protein-protein interaction

**DOI:** 10.1101/2024.06.03.596235

**Authors:** Ian Hope, Martin E. M. Noble, Michael J. Waring, E. M. Martin Noble, Jane A. Endicott, Natalie J. Tatum

**Affiliations:** Newcastle University Centre for Cancer, Translational and Clinical Research Institute, Newcastle University, Paul O’Gorman Building, Newcastle upon Tyne, NE2 4HH, UK; Cancer Research Horizons Therapeutic Innovation, Newcastle Drug Discovery Group, Translational and Clinical Research Institute, Newcastle University, Paul O’Gorman Building, Newcastle upon Tyne, NE2 4HH, UK; Chemistry, School of Natural and Environmental Sciences, Newcastle University, Bedson Building, Newcastle upon Tyne, NE1 7RU, UK

**Keywords:** cyclin, protein kinase, regulation, X-ray crystallography, inhibitor design, cell cycle, protein kinase, protein-protein interactions, structure guided inhibitor design

## Abstract

Protein-protein interaction sites (PPIs) are potentially more selective therapeutic binding sites than protein substrate binding sites. PPIs include distinct regions frequently called “hotspots,” sites of key amino acid interactions. Prospective identification of these hotspots through X-ray crystallographic screening could assist in the identification of separation of function mutants for experimental validation, enhance confidence in AI-generated multiprotein complex predictions and accelerate development of selective chemical probes. To explore these applications, we utilize the FragLite library to examine the binding surfaces of CDK2-cyclin A. The many protein- and peptide-CDK2-cyclin A complexes that have been structurally characterised make this complex an appropriate test case. We show that FragLites comprehensively map both known sites of protein-protein interaction on CDK2-cyclin A and identify a possible uncharacterised site, providing a structural method toward directing mechanistic studies and providing starting points for chemical probe design.

## Introduction

Cyclin-dependent kinases (CDKs) are a subgroup of serine/threonine protein kinases that are activated by cognate cyclin binding.^1–3^ CDK-cyclin complexes are crucial both for orchestrating transitions through the cell cycle, and for regulating transcription. As examples, the cell cycle CDKs, CDK1, 2, 3, 4 and 6 bind to group I cyclins, namely cyclins A (CDKs 1 and 2), B (CDK1), D (CDKs 4 and 6), and E (CDK2). The CDK subfamily that regulates gene transcription are similarly distinguished and have preferred cyclin partners, CDKs 7, 8, 9 and 12 pairing with cyclins H, C, T and K respectively.

Cyclins activate their cognate CDKs by remodelling the CDK active site, an allosteric event that is accompanied by CDK phosphorylation within the activation segment, which in CDK2 comprises residues 145-172 between the conserved DFG and APE motifs (single letter amino acid code).

CDK2 is phosphorylated on Thr160 and partners with cyclin A2 to control cell cycle progression through S-phase into G2.^4,5^ Both CDK2 and cyclin A bind to CDK2 substrates and proteins that regulate CDK2 activity. The CDK2 C-terminal lobe contains the GDSEID sequence within a loop linking the αF and αG helices (residues 205-210) at which interactions with CDK-associated protein 1 (CKS1)^6^, kinase associated phosphatase (KAP)^7^ and M-phase inducer phosphatase 1 (CDC25A)^8^ have been structurally characterised. The cyclin A recruitment site on the N-terminal cyclin box fold (N-CBF), which is also conserved in B, D and E cyclins, recognises CDK substrates and protein inhibitors that contain the RXL motif. ^9,10^

Protein-protein interactions (PPIs) commonly include hotspot regions, small finite areas of the interactome with a concentrated number of key residues that contact the partner protein and make a significant contribution to binding free energy.^11,12^ PPIs can be formed by two structured interfaces, and also, as exemplified by CDK-cyclin complexes, through short linear motifs (SLiMs).^13,14^ Around 2-12 amino acids in length, SLiMs have a defined sequence within intrinsically disordered regions of the proteome and, in the case of CDK-cyclin complexes, dock to sites on cyclins with low affinity, allowing site- and temporally-specific regulation of CDK-cyclin signalling cascades.^15–19^

The first cyclin binding SLiM to be characterised was the RXL motif present in many cell cycle CDK-cyclin substrates including E2F, pRb, ORC1 and p27.^9,10,18,20–23^ The RXL recruitment site on the cell cycle cyclin subfamily is a characteristic hydrophobic patch, defined on cyclin A2 by the MRAIL sequence on helix α1 in the N-CBF.^22,24^ Positional variation in the RXL motif and surrounding residues can tune CDK substrate affinity. ^1,10,21,25,26^

Profiling of CDK substrates in yeast has helped to identify SLiMs that contribute to substrate recognition at each stage of the cell cycle.^17^ Recent structural characterisation of the human separase-Cdk1-cyclin B1-CKS1 complex revealed how multiple SLiM docking interactions may collectively achieve CDK-substrate selectivity.^27^ Mutational studies to SLiM sequences have shown these recognition elements to be key contributors to PPIs that enable CDKs to interact with various binding partners, including cyclins, CDK inhibitors (CKIs), and other regulatory proteins.^14,17,27–29^ Recent structural studies on the transcriptional CDK-cyclin family have identified additional sites of protein-protein interaction within the C-terminal cyclin box fold (C-CBF) that also exploit both structured interfaces and SLiMs.^30–32^ It is now apparent that in a CDK-cyclin module-dependent manner, a significant fraction of the cyclin surface mediates protein interactions.

A large number of potent and selective ATP-competitive inhibitors have been designed that can be used as probes to study CDK function.^33–38^ However, they suffer from being agnostic as to peptide substrate identity and their impact on CDK-protein interactions is difficult to deconvolute.

Perturbing interactions made by CDK-cyclin complexes, particularly at SLiM sites, presents an opportunity to design more selective and discriminative probes of CDK function, which would bypass the orthosteric ATP binding site and provide specificity to both CDK activation states and particular signalling pathways.^29^ Taking cyclin A as an example, truncation of RXL-containing substrates into peptidomimetic fragments has been explored both to optimise the cyclin binding motif to inhibit CDK-substrate interactions ^9,22,39–43^ and to provide selective cyclin A binding moieties to enable early PROTAC design.^44^ The ability of small molecule CDK inhibitors to perturb PPIs has recently been highlighted by the characterisation of inhibitors I-125A and I-198 that bind at the CDK2-cyclin E1 interface to remodel both a loop present in cyclin E1 and the activation segment of CDK2 to stabilise a structure that is not capable of peptide substrate recognition and catalysis.^45^ The exquisite selectivity of these compounds for CDK2-cyclin E1 highlights the potential of targeting allosteric sites for drug development and the importance of interface agnostic methods to uncover new potential protein interaction sites.

FragLites are a 31-member library of small, halogenated fragments, which specifically and selectively map binding sites on proteins as effectively as a full fragment screen.^46,47^ The FragLite mapping of monomeric CDK2 in its inactive conformation identified fragment clusters that overlapped known substrate and protein-protein interaction sites, rationalising the use of the library as an effective tool for preferentially mapping PPI hotspots. We have used FragLite technology to map the CDK2-cyclin A complex, probing, for the first time, the interactome of cyclin A and allowing comparison to known features of CDK2-cyclin A-containing complexes. This analysis characterises known CDK2-cyclin A protein binding sites and identifies a potential protein binding site at the CDK-cyclin A interface. It also provides a chemically rich map of the CDK2 and cyclin A protein surfaces as a starting point for chemical probe design and to aid mutation site selection to distinguish functions.

## Results

Thirty-one FragLites (**Table S1**) were soaked individually into crystals of the CDK2-cyclin A complex and subjected to X-ray diffraction (**Figure 1A-1D, and Table S2**). A characteristic of the FragLites is the ability to exploit the anomalous scattering of either the bromine or iodine atom incorporated into their structures to unambiguously identify binding events (**Figure S1A**). Ligand binding was determined using an overlay of the anomalous log-likelihood gain (LLG) map with the 2Fo-Fc electron density map and Fo-Fc difference map and searching for difference anomalous peaks greater than 4 standard deviations above the mean value. Thirty crystals generated diffraction patterns, with only FL21 compromising crystal diffraction (**Table S2**). Further inspection of ligand binding revealed only two datasets with no detectable binding event (FL15 and FL25) delivering a hit rate of 90%. This hit rate exceeds that observed for the monomeric CDK2 complex (29 %)^47^, and for the bromodomains of ATAD2 and BRD4 (39 % and 65%, respectively)^46^. Most FragLites bound at more than one site, and as the CDK2-cyclin A complex crystallises as a dimer of dimers in the asymmetric unit, FragLite binding was often detected in both copies.

**Figure 1.**
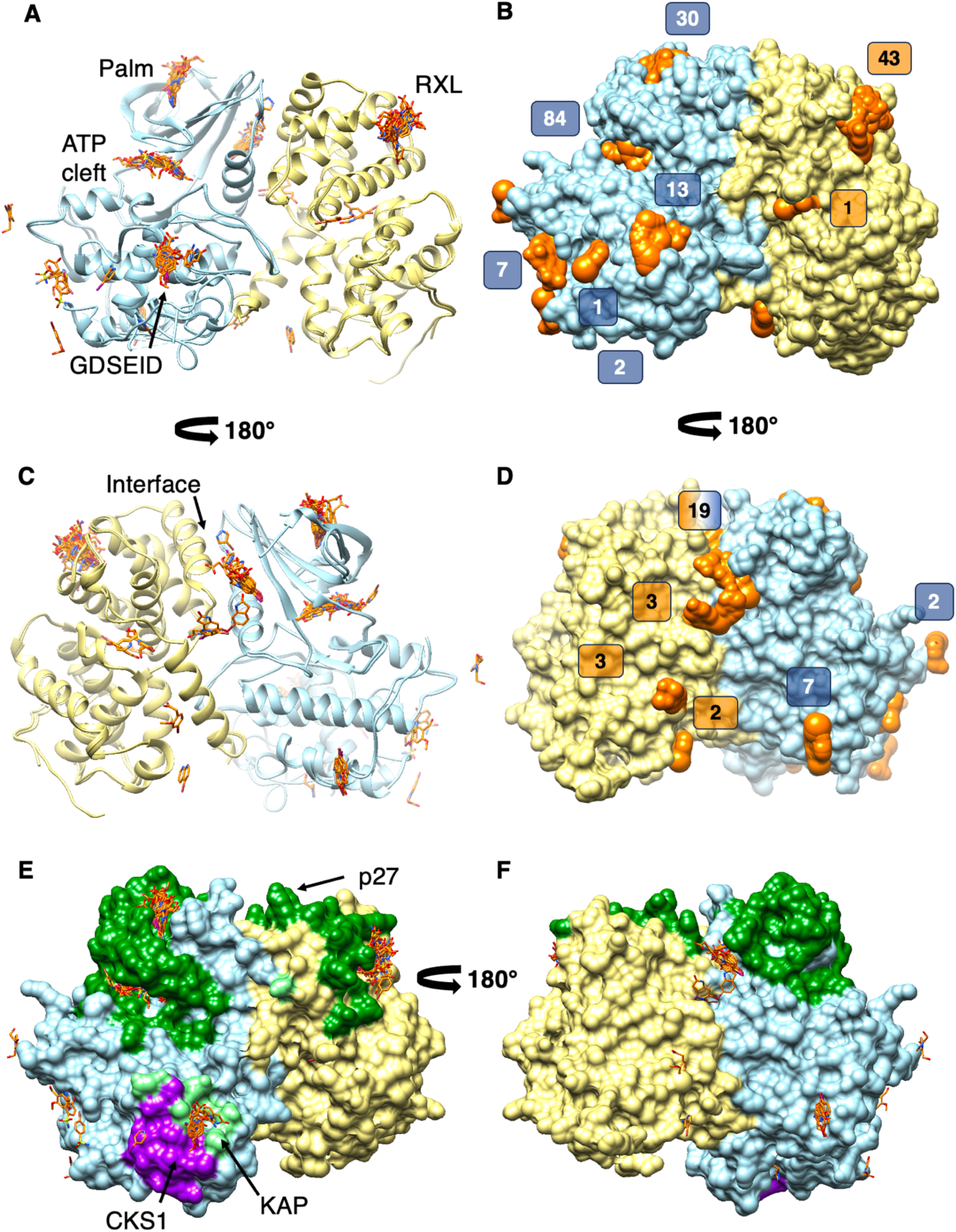
The CDK2-cyclin A FragLite map. **(A, C)** Superposition of the CDK2-cyclin A dimer of dimers collates all hits against the active CDK2-cyclin A complex. There are significant clusters of FragLite binding in the Palm and ATP-binding cleft of CDK2, and at the RXL site of cyclin A (30, 84 and 43 binding events respectively). Other notable FragLite binding sites are the CDK2 GDSEID motif (13 binding events) and a site at the CDK-cyclin A interface (19 events). CDK2 and cyclin A are shown in pale blue and yellow ribbon respectively with FragLite binders in orange sticks coloured by heteroatom; ribbon shown is of CDK2-cyclin A-FL1 complex. **(B, D)** Molecular surfaces of CDK2-cyclin A (from CDK2-cyclin A-FL1 complex) and FragLites demonstrate the space filled by fragment coverage across the complex surface. The total number of binding events is 217, numbered here by cluster on CDK2-cyclin A. (**E, F**) FragLites map known CDK2-cyclin A-protein interaction sites. CDK2 and cyclin A residues that contact CKS1, KAP and p27 extracted from PDB entries 1BUH (CKS1), 1FQ1 (KAP) and 1JST (p27) are coloured magenta, light green and dark green respectively on the molecular surface of the CDK2-cyclin A-FL1 complex. The FragLite map is included for comparison. The view in panels A, B and E is equivalent as is the view selected in panels C, D and F.

The consolidated map (**Figure 1A-1D**) shows that FragLite binding is strongly but not exclusively clustered in the active site of CDK2 and at the RXL recruitment site of cyclin A (84 and 43 binding events respectively, **Figure 1B**). FragLite binding does not change the architecture of the CDK2 ATP binding site, and the structures are locally indistinguishable from the structure of CDK2-cyclin A bound to ATP (PDB 1FIN)^48^. The more extensive FragLite coverage with respect to the FragLite map of monomeric CDK2 (**Figure S1B**) arises at least in part from the crystallographic dimer of dimers, wherein fragments bind to one or both copies of CDK2 or cyclin A within the asymmetric unit (**Figure S2A**). The consolidated map (**Figure S2B**) highlights that these events are not technical replicates. Each dimer presents a distinct crystal environment through varying crystal contacts (**Figure S2C-S2D**) and the binding modes of each FragLite can differ with respect to each dimer.

The ATP binding site and “Palm site” (**Figure 1A**) formed within the CDK2 N-terminal twisted β-sheet were identified by the monomeric CDK2 FragLite map.^29,47^ However, the environment within the CDK2-cyclin A crystals leads to richer engagement at these sites. The CDK2-cyclin A map also highlights the CDK2 C-terminal lobe around the GDSEID sequence that precedes helix αG (13 events) (**Figure 1A**), the cyclin A RXL recruitment site (43 events) (**Figure 1A**), and a site formed at the interface between CDK2 and cyclin A (19 events) (**Figure 1C**). As illustrated in **Figure 1E** and **1F**, FragLites map known sites of protein interactions.

## Discussion

### FragLite engagement sites on CDK2 p27

p27 is an intrinsically disordered protein that within its N-terminal kinase interaction domain (KID) features sequence motifs that engage with the cyclin recruitment site, the CDK2 N-terminal lobe and the ATP binding cleft.^49,50^ Though the crystal structure of the ternary CDK2-cyclin A-p27 complex suggests p27 ordering on binding,^22^ in solution methods are consistent with a more dynamic ternary assembly.^49–51^ p27 interactions with CDK2 engage the CDK2 N-terminal lobe within the Palm site before displacing the edge strand of the N-terminal β-sheet to enter the CDK2 active site.^22^ Collectively these interactions prevent peptide substrate and ATP binding, and re-arrange conserved residues required for catalysis.

Fifteen FragLites bind at the Palm site (**Figure 2A**). All bury their bromine substituent and aromatic core, consistent with their ability to emulate aromatic sidechain binding. The most populous binding pose (Palm site mode 1) is approached from the Palm’s opening and is exemplified by FL1/2 (**Figure 2B**). The hydrogen bonding properties of the FragLite doublet drives a preference in FragLite orientation. A subset of FragLites which share the same relative donor/acceptor dispositions (FL1, 2, 4, 5, 18, 22, 23) form hydrogen bonds with the backbone carbonyl oxygens of Met1 (3.4 Å) and Phe4 (2.8 Å). FL1 and FL2 exhibit a distinct second binding pose (mode 2) which buries their imidazole core within the Palm site and engages in a side-on π-stacking interaction with Tyr19 while forming hydrogen bonds with the backbone carbonyl oxygen of Val17 (2.8 Å).

**Figure 2.**
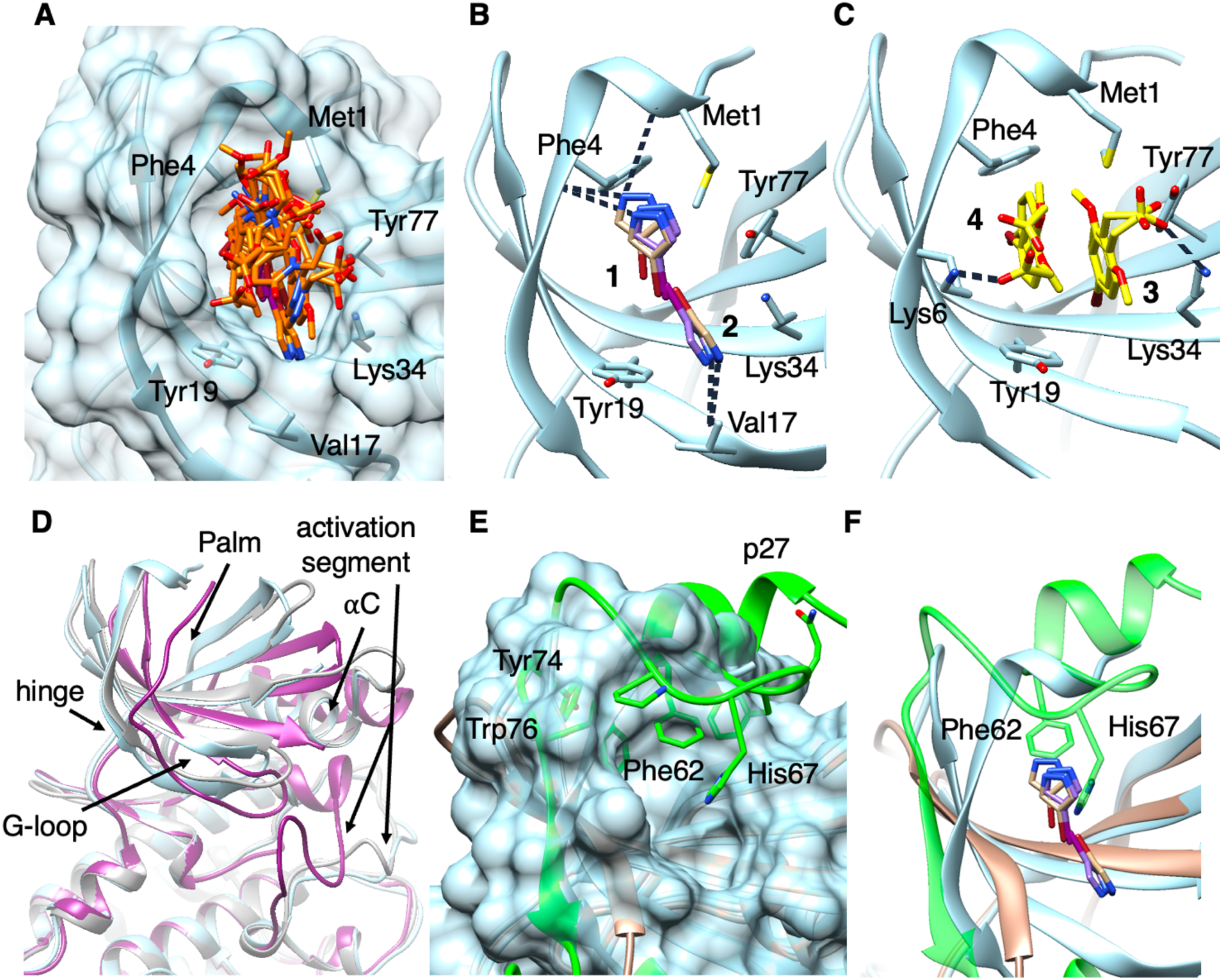
FragLite engagement in the CDK2 Palm. **(A)** The cluster of FragLites bound to the Palm of CDK2-cyclin A with molecular surface of the CDK2-cyclin A-FL1 complex shown. FragLites are shown in orange sticks coloured by heteroatom. **(B)** FL1 (tan carbons) and FL2 (purple carbons) primarily form interactions with backbone donors and acceptors; the halogen (brown for Br, purple for I) is buried in the centre of the Palm to orient the donor/acceptors of the FragLite toward hydrogen bonding partners such as Met1 (3.4 Å), Phe4 (2.8 Å) and Val17 (2.8 Å). **(C)** FL31 molecules stack within the Palm in modes 3 and 4, forming hydrogen bonds to Lys6 (2.3 Å) and Lys34 (3.0 Å) side-chains. **(D)** The Palm conformation in the FragLite-bound CDK2-cyclin A complex (blue ribbon) is similar to the *apo* complex exemplified by PDB 1FIN (grey ribbon), though notably relaxed compared to the FL1-bound structure of monomeric CDK2 (PDB: 6Q3C, magenta ribbon). Key secondary structure elements of CDK2 including the activation segment are labelled. **(E)** The Palm is a site of p27 binding (PDB: 1JSU, CDK2 in bronze, p27 in green, FragLite-bound CDK2-cyclin A-FL1 surface in blue). The Palm is considerably more open in the inactive p27-bound complex than the active state, to accommodate burial of the aromatic p27 residues, chiefly Phe62 and His67. Tyr74 is exposed to solvent. **(F)** The positioning of FL1 and FL2 mirror the burial of Phe62 and His67 from p27 (overlaid from PDB 1JSU, p27 in green, CDK2 in bronze).

Adopting an alternative binding mode, a second subset (FragLites 7, 19, and 31) frequently direct the acceptor groups towards CDK2 Tyr77 to form hydrogen bonds with the Tyr77 hydroxyl group (Palm site mode 3, exemplified by FL31, **Figure 2C**). FL31 also adopts two binding poses. In one copy FL31, in addition to capturing a hydrogen bond with Tyr77, also makes additional ionic interactions through its carboxylate group to Lys34 (3.0 Å). In a second orientation, only observed with FL31 (mode 4), it captures a T-shaped interaction with the phenol sidechain of Tyr19 and an ionic interaction between the carboxylate group with Lys6 (2.3 Å) (**Figure 2C**). Notably, the two bound copies of FL31 π-stack against each other.

The conformation of the Palm site is highly malleable and dependent on the activation state of CDK2 (**Figure 2D**). In the inactive monomeric CDK2 structure, in addition to the ⍺C helix being pushed out and the activation segment folded in, the Palm adopts a tight closed conformation (**Figure 2D**, magenta fold.^47^ Notably, only one FragLite (FL31) was observed to bind to the monomeric CDK2 Palm site. Within the active CDK2-cyclin A complex, the site is more open (**Figure 2D**, grey fold^52^ and FL1-bound, blue) it is the second most populous FragLite binding site on CDK2 (**Figure 1A, B**). The Palm site is a docking pocket for p27 aromatic residues 62-76 (**Figure 2E**) which includes Tyr74, the residue that plays a pivotal role as a phosphoswitch for CDK engagement.^29,49^

FragLite interactions within the Palm site are reminiscent of those displayed by p27. Although the re-arrangements to the CDK2 structure in this region that accompany p27 binding do not occur on FragLite binding (compare **Figure 2D**), FragLites do occupy an important region in which p27 binds several of its highly conserved aromatic residues (Phe62, Phe64, His67, **Figure 2E**).^22,29^ Of these residues, Phe62 and His67 are most proximal to FragLite binding (illustrated for FL1 and FL2 binding in **Figure 2F**). FragLites make a hydrogen bond with the sidechain of CDK2 Tyr77, resembling as illustrated for FL31 and FL1/2 (**Figure 2E** and **2F** respectively) that which CDK2 makes with p27 His67. Emphasizing the importance of p27 residues Phe62 and Phe64 to p27 binding to CDK2-cyclin A, alanine substitutions abrogate binding.^53–55^

FragLite binding to the ATP binding site recapitulates the interactions between CDK2 and p27 and CDK2 and ATP. FragLite binding in the ATP-binding site of CDK2 can be categorised into two binding modes. The major binding mode (ATP site mode 1) represents two-thirds of the events in this pocket and is exemplified by FL1 and FL2 (carbons coloured tan and purple, **Figure S3A**), in which a pyrazole nitrogen forms a hydrogen bond with either the backbone amide of Leu83 (2.8 Å) or the carbonyl of Glu81 (2.9 Å) within the CDK2 hinge sequence linking the N- and C-terminal lobes. FragLites adopting mode 1 overlay the ATP adenine moiety. The remaining events assume an alternate pose (ATP site mode 2) in which the halogen atom is directed away from the inner ATP site cleft toward Glu81 and Leu83. Mode 2 is exemplified by FL8 (**Figure S3B**), demonstrating this position allows FragLite hydrogen bonding motifs to interact with the polar pocket formed by Lys33, Lys89, Glu51 and Asp145. In this binding mode the FragLites make no hydrogen bond or halogen bond interactions with the CDK2 backbone at the hinge sequence; instead FL8 forms hydrogen bonds to the Lys33 sidechain and Asp145 backbone. p27 residues Phe87 and Tyr88 also bind deep within the ATP cleft (**Figure S3C**) and FragLites mimic the positions of these key p27 residues in both binding modes, as exemplified by FL1, FL8 and FL18 (carbons coloured tan, sienna and pale yellow respectively, **Figure S3D**).

### The CDK2 GDSEID motif

The GDSEID motif preceding the CDK2 helix αG on the C-terminal lobe of CDK2 binds KAP (PDB 1FQ1),^7^ CKS1 (PDB 1BUH)^6^ and CDC25A (PDB 8ROZ).^8^ The C-terminal helix of KAP, specifically Ile183 and Tyr186 interact with a non-polar recognition site on CDK2 composed of the GDSEID motif and residues 235-237 (**Figure 3A**). CKS1 also presents a complementary apolar/aromatic residue pairing in the same region, such that CKS1 His60 and KAP Ile183, and CKS1 Leu68 and KAP Tyr186 respectively resemble each other in binding mode (**Figure 3B**). CDC25A forms more extensive hydrogen bonding interactions between its C-terminal helix and the CDK2 αG helix (**Figure 3C**), where Asp206 and Asp210 of the GDSEID form hydrogen bonds CDC25A to Arg450 and Tyr455 respectively. This extended CDC25A interface at the GDSEID binding region supports recognition of CDK2 as a substrate by CDC25A.

**Figure 3.**
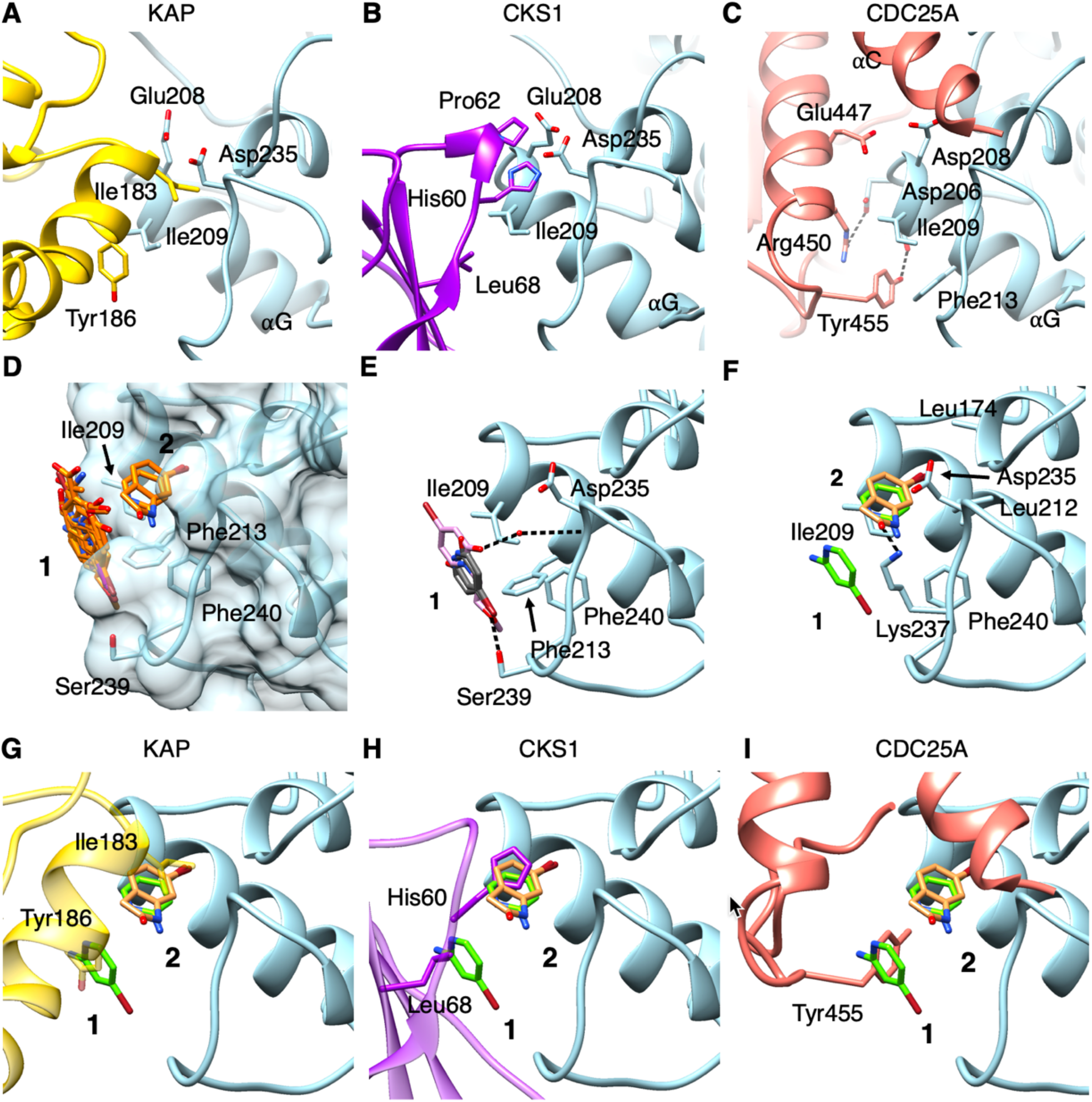
FragLite engagement at the GDSEID site. **(A)** KAP (gold ribbon) interacts with CDK2 (blue ribbon) via key residues Ile183 and Tyr186 (PDB:1FQ1). Glu208 is the first residue of the GSEID sequence at the N-terminal end of the ⍺G helix. **(B)** CKS1 (purple ribbon) also interacts with the N-terminal end of the CDK2 αG helix through His60 and burial of Leu68. **(C)** CDC25A (salmon ribbon) forms extensive helix-helix contacts along the CDK2 αG helix, including hydrogen bonds from Tyr455 and Arg450 to Asp 206 (3.4 Å) and Asp210 (2.4 Å) of the GDSEID motif respectively. **(D)** FragLites bind at the GDSEID site in two clusters, each representing a separate mode where mode 1 orients parallel to the CDK2 surface and mode 2 buries its halogen towards the GDSEID helix. **(E)** Mode 1 is exemplified by FL6 (grey carbons), which forms a hydrogen bond to Ser239 (3.5 Å), orienting the Br atom, and FL29 (pink carbons) which orients the methoxy substituent down to engage with Ser239 (2.9 Å) and forming a hydrogen bond to Asp235 via a water molecule (2.8 Å, HOH-Asp235 3.2 Å). **(F)** Mode 2 is exemplified by FL10 (sandy brown carbons) and the upper event of FL4 (green carbons), which are each sandwiched between Ile209 and Asp235. Each forms a hydrogen bond with Lys237 (FL4, 3.5 Å; FL10, 2.3 Å). **(G)** Modes 1 and 2 mimic KAP binding. KAP shown in transparent gold ribbon with key residues Tyr186 (mimicked by mode 1) and Ile183 (mimicked by mode 2) shown for direct comparison with panel F. **(H)** Modes 1 and 2 mimic CKS1 binding (purple ribbon), where mode 1 mimics key residue Leu68 and mode 2 mimics His60. **(I)** Modes 1 and 2 mimic to a lesser extent CDC25A engagement (salmon ribbon), which is dominated by the extensive helix-helix interface rather than a hotspot-like engagement.

Ten FragLites bind adjacent to the αG-helix in two distinct clusters (**Figure 3D**). In the first cluster (GDSEID site, mode 1), the aromatic FragLite core caps the face of Ile209 and the bromine atom is typically directed toward a main chain hydrophobic pocket created by Ile209, Phe213 and Phe240 with evidence of a halogen bond to the sidechain of Ser239 (3.3 Å as illustrated for FL6 Br, **Figure 3E**, grey carbons). In contrast to the majority of FragLites adopting this binding pose, a different orientation of the bromine atom is observed for FL29 (**Figure 3E**, pink carbons). The FL29 aromatic core overlaps with the other FragLites and interacts with Ile209, but in contrast it is its methoxy tail and not its halogen atom that interacts with Ser239 through a hydrogen bond (3.5 Å). The pyridone oxygen creates a hydrogen bonding network through coordination with nearby water molecules to the main chain carbonyl of Asp235.

The second cluster of FragLites (GDSEID site, mode 2) is exemplified by FL4 and FL10 (carbon atoms in green and sandy brown respectively, **Figure 3F**). One copy of FL4 binds in mode 1, but in the second, the FL4 aminopyridine core overlays the FL10 indolinone ring so that they make equivalent interactions with neighbouring residues (mode 2). Both FragLites bury their bromine in a pocket composed of residues Leu174 and Leu212, Ile209 and Phe240. Binding is further supported by an anion π-complex between Asp235 and the electron deficient core, and the formation of a hydrogen bond (2.3 Å) with Lys237.

Each of these two modes emulates KAP/CKS1/CDC25A engagement (**Figures 3G, 3H** and **3I** respectively), exemplified by FL4 and FL10. Mode 2 emulates the CKS1 His60/KAP Ile183 interaction whereas mode 1 imitates the CKS1 Leu68/KAP Tyr186 engagement. Notably site-directed mutagenesis of CKS1 residues (Tyr57Ser, Met58Ala, His60Asp, Glu63Lys)^6^ spanning both FragLite pockets has been shown to disrupt the CDK-CKS1 interaction, providing evidence for the significance of this docking site. As a more extended interaction surface rather than a hotspot-site, residues on CDC25A show no direct overlap with FragLite events at the GDSEID site, though FL4 emulates somewhat the Tyr455 protrusion toward Ile209. Collectively, these findings support a dual pocket enhanced by specific aromatic/hydrophobic pairings, serving as a shared docking site for regulatory proteins. In emulating CDK2-protein interactions the FragLites identify CDK2 residues to guide the generation of separation of function mutations to selectively distinguish CDK2 substrate selection (CKS1 binding) and regulation (KAP vs CDC25A binding) for mechanistic studies.

### Sites on cyclin A RXL site

Our screen identified a total of five FragLite binding sites on cyclin A (**Figure 1B, D**). Nineteen FragLites bound to the RXL recruitment site occupying both copies of CDK2-cyclin A within the asymmetric unit resulting in a total of 43 binding events (**Figure 4A**). Two binding modes are evident in the recruitment site (**Figure 4B**). In all orientations, FragLites orient their bromine substituents into the recruitment site that characterises the RXL recruitment site. In binding mode 1, FragLites direct their halogen substituent to form a halogen-π interaction with Trp217 and are encapsulated by Ile213 and Gln254. For the first cyclin A copy in the crystal, FragLites (as exemplified by the binding of FL4) bind face-to-face in a 2:1 FragLite:cyclin A ratio; In the second copy, the second FragLite binding site is occupied by the sidechain of Leu96 from a symmetry-related CDK2 molecule (not shown). In RXL site binding mode 2, FragLites are sandwiched between members of the MRAIL helix; Met210, Ile213, Leu214 and Leu253 in helix α3, exemplified by FL17. Two FragLite compounds (FL23 and FL27) also adopted a third binding mode, engaged in a π-π stacking interaction with Trp217. Specifically, FL23 (carbon atoms in blue, **Figure 4C**) is within hydrogen bonding distance to the carbonyl backbone of Ile281 (2.9 Å) and the sidechains of Gln254 (3.0 Å), in contrast to FL27 (not shown) whose interactions with these residues are mediated by neighbouring water molecules.

**Figure 4.**
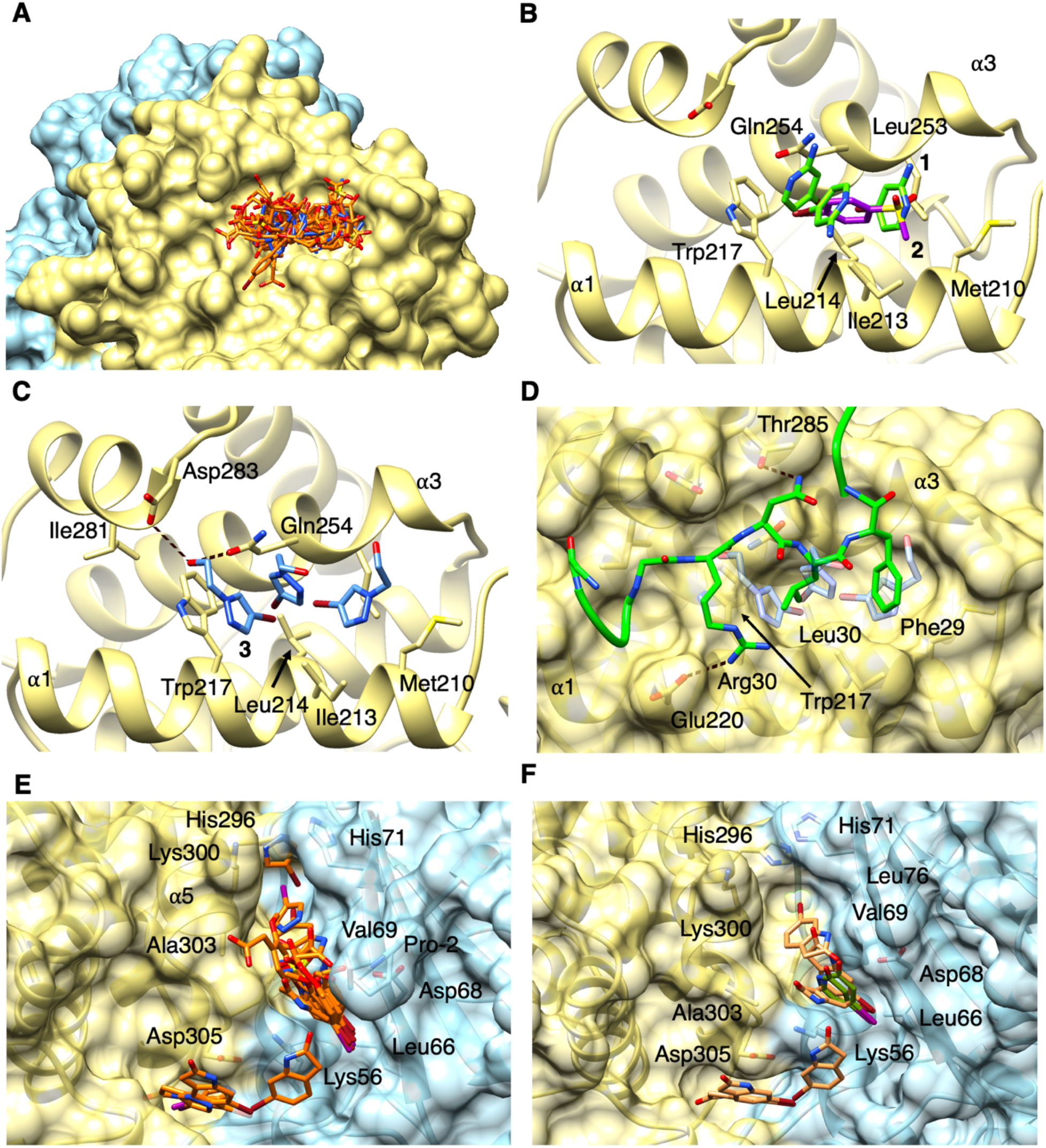
FragLite engagement on cyclin A. **(A)** A total of 43 binding events occur in the recruitment site which binds RXL-containing substrates. **(B)** FragLites adopt one of two binding modes in the recruitment site, either perpendicular burial of the halogen as exemplified by FL4 (mode 1, green carbons), or a parallel binding as exemplified by FL17 (mode 2, purple carbons). Met210 begins the MRAIL motif characterising the recruitment site formed by helices ⍺1 and ⍺3. **(C)** FL23 events (blue carbons) exhibit stacking with key recruitment site residue Trp217 and forms hydrogen bonds with the Ile281 backbone (2.9 Å). **(D)** The RXLF motif of p27 (PDB: 1JSU, shown in green ribbon and carbons) buries Leu30 and Phe29 into the recruitment site of cyclin A, while forming hydrogen bonds to Glu220 and Thr285. Overlay with transparent FL23 demonstrates the mimicry of FragLites to these interaction motifs. **(E)** A site on the interface between CDK2 and cyclin A (CDK2-cyclin A-FL2 surface and side chains shown). FragLites map this site formed by the cyclin A N-CBF residues His296, Lys300, Ala303 and Asp305, and the N-lobe CDK2 residues His71, Val69, Asp68, Leu66, Lys56 and the Pro at position -2 from the protease cleavage site. **(F)** FL7 (dark green carbons) and FL10 (sandy brown carbons) demonstrate the main binding mode wherein the halogen buries into a recruitment site around Leu66. FL10 molecules effectively map the contours of this interaction site from the His296:His71 interaction down to a trench beneath Asp305.

FragLites emulate features that contribute to p27 binding to the recruitment site (**Figure 4D**). In the CDK2-cyclin A-p27 structure, p27 orients Leu32 and Phe33 of its RXLF motif into the cyclin A surface.^22^ As exemplified by FL23 there is a close overlap of the FragLite cores with the sidechains of this hydrophobic pair of residues and FL23 uniquely overlaps the backbone carbonyl oxygen of p27 Arg30 of the RXL motif, mimicking this interaction via the methoxy oxygen acceptor.

### Potential to identify novel interaction sites

A richly populated site (19 binding events) at the CDK2-cyclin A interface was observed (**Figure 1**). The site is formed by cyclin A residues His296, Lys300, Ala303 and Asp305, and CDK2 residues Lys56, Leu66, Asp68, Val69, His71, and Leu76. Within this extended site, FragLites exhibit a common orientation, featuring a shared cation π-interaction between the aromatic core of each FragLite and the sidechain of CDK2 Lys56 (**Figure 4E** and **4F**). The main binding mode within the cleft is exemplified by FL7 (**Figure 4F**, dark green carbons), wherein the halogen substituent is directed toward the hydrophobic groove formed by CDK residues Lys56, Leu66, Asp68, and Val69. Additionally, the hydrogen bonding substituents of FragLites are commonly found within hydrogen bonding distance to CDK2 residues Asp68 and His71. In contrast, FL10 (carbon atoms in sandy brown) embeds the halogen into the hydrophobic pocket (formed by cyclin A residues Phe304 and His296, and CDK2 residues Leu76 and His71) in multiple orientations through the interface groove, framing Lys56 (**Figure 4F**). Evaluated together, the events map the extent of the interface site surface.

This site is not a known protein interaction site. The six CDK2 and four cyclin A2 residues that create it are highly conserved between mammalian as well as *Drosophila melanogaster* and *Caenorhabditis elegans* CDK2 proteins and close human homologue CDK1; and between mammalian cyclin A2 as well as *Xenopus laevis* cyclin A2 and close human homologue cyclin B1. This pattern of high conservation even among more distant genera suggests a role of as-yet-undefined functional significance. This newly identified site will be the focus of future studies.

Occlusion of protein interaction sites by crystal lattice contacts is a limitation of employing crystallographic screens to identify protein-protein interaction sites. Identifying alternative crystal forms for the protein of interest is demonstrated by this study to enrich fragment maps and provides greater confidence in the identification of potential protein interaction sites; access to such alternative forms may be achieved by varying construct design or crystallisation conditions.^56^ A comparison of the CDK2 and CDK2-cyclin A FragLite maps shows the GDSEID site on CDK2 was bound by FragLites in crystals of CDK2-cyclin A but not in crystals of monomeric CDK2, where it was occluded by a lattice contact. Similarly, FragLite screening failed to identify the SKP2-specific binding site within the cyclin A N-CBF.^57^ The cyclin A residues that interact with SKP2 (specifically residues 244-249 that include the SSMS motif) are partially occluded by the β2-β3 turn of the

CDK2 Palm in the second copy of CDK2 present in the asymmetric unit. These examples demonstrate crystal contacts, themselves a protein:protein interaction, should not be discounted as uninformative. Additionally, we observe some sites occupied by a smaller (≤ 3) number of FragLites which are typically supported by crystal contacts (**Figure S2C** and **S2D**). The utility of FragLites to bind in richness to sites of functional relevance, corroborated by mutational studies, therefore allows holistic prioritisation of highly occupied sites when evaluating a new target.

## Significance

This study highlights that while singleton fragments may opportunistically bind to available surfaces and sites, richness of fragment binding reflects more significant functional hotspots. By their design, FragLites can map sites effectively and efficiently with a small number of chemical entities. Known sites of CDK2-cyclin A protein interaction are the most populous sites, and it is hotspots (rather than extended interaction surfaces) that bind FragLites. Through comparison to docking studies, we have demonstrated the ability of these fragments to preferentially identify sites of residue-specific interaction. In identifying a highly populated but undocumented site on the CDK2-cyclin A interface, we suggest FragLites may have utility in prospective mapping.

Furthermore, we note that these maps can enhance the confidence placed in AI-generated structural models of multiprotein complexes. Surface mapping by crystallographic fragment screening has to date been primarily explored as a drug discovery method to identify chemical starting points to develop allosteric inhibitors.^58,59^ Our FragLite analysis underscores the potential of FragLites both as tools to aid development of chemical probes and as an approach to identify potential separation of function mutants to investigate target biology.

## Star * Methods

## Contact for Reagent and Resource Sharing

Further information and requests for resources and reagents should be directed to the corresponding author.

## Method Details

### Cloning, expression and purification of CDK2-cyclin A

Generation of CDK2-cyclin A has been reported previously.^60^ Briefly, co-expression of CDK2 with *S. cerevisiae* CAK1 generates CDK2 stoichiometrically phosphorylated on Thr160 within the activation segment (hereafter referred to as CDK2). Full-length human CDK2 (P24941) and *S. cerevisiae* CAK1 (P43568) were co-expressed as GST fusions from independent mRNAs from the pGEX-6P-1 vector backbone (Cytiva). Bovine cyclin A2 (P30274) residues 170-430 is expressed independently from a second vector based on the pET-21d(+) backbone (Novagen). This sequence is equivalent to human cyclin A2 (P20248) residues 172-432. The human numbering has been used throughout. As a result of cloning artefacts, the CDK2 sequence is preceded by the sequence GPGS at the N-terminus and bovine cyclin A2 contains a non-cleavable His6 tag at the C-terminus. The two plasmids were independently transformed into *Escherichia coli* BL21 (DE3) pLysS competent cells. For each construct, single colonies were selected to initiate a 10 ml overnight starter culture which was then transferred to 1L 2YT autoinduction media (Formedium). Cells were subsequently grown at 30 °C for 16-20 hr prior to harvesting by centrifugation (4000 xg). Cell pellets expressing bovine cyclin A2 and co-expressed CDK2 and CAK1 were each resuspended in mHBS (200 mM NaCl, 40 mM HEPES pH 7.5, 1 mM TCEP). Suspensions were combined in a ratio of 2:1 by cell culture volume, cyclin A2: CDK2-CAK1 and then supplemented with 2 μg mL^-1^ DNase I, 10 μg mL^-1^ RNase A, 25 μg mL^-1^ lysozyme and 5 mM MgCl_2_, and lysed by sonication (5 min total, pulsed 20 s on and 40 s off, 4°C, (Sonics vibra cell sonicator at 30 % amplitude (VCX 500 / VCX 750, Sonic & Materials, CT, USA)). Lysates were then clarified by centrifugation (100,000 xg, 60 min, 4 °C, JA 30.50 rotor, Avanti JXN-26 centrifuge (Beckman)) before filtering the supernatant through 0.45 μm syringe filters. Recovered lysates were bound to GST resin (Cytiva; pre-equilibrated with mHBS) via batch binding before washing to baseline and then eluting the complex in 200 mM NaCl, 40 mM HEPES pH 7.5, 1 mM TCEP, 20 mM reduced glutathione, pH checked to 7.5. Eluted GSTCDK2-cyclin A complex was then cleaved overnight (16 hr) at 4 °C using 1:50 mass:mass of HRV 3C protease:GSTCDK2-cyclin A. Cleaved product was filtered (0.2 μm, Minisart®, Sartorius, Germany) and subject to size-exclusion chromatography (SEC) using an s75 preparative grade Superdex® HiLoad® column (Cytiva) pre-equilibrated in mHBS at 4°C. Peak fractions were verified using SDS-PAGE before pooling and removing trace GST by subtractive glutathione-affinity chromatography using a 5 mL GSTrap FF column (Cytiva) equilibrated in mHBS. Fractions containing CDK2-cyclin A were pooled and concentrated to 10 mg mL^-1^ and stored at -80 °C. Protein concentrations were measured at OD_280 nm_ using a Nanodrop Spectrophotometer.

### Crystallisation, X-ray fragment screening and structure determination of CDK2-cyclin A with fragments

Crystallisation of CDK2-cyclin A was performed using a vapour diffusion sitting drop method in MRC 2 Lens crystallisation plates (SWISSCI) using a crystallization matrix (0.6-0.8 M KCl, 0.9–1.2 M (NH_4_)_2_SO_4_, and 100 mM HEPES pH 7.0) prepared in house using a Biomek liquid handler instrument (Beckman Coulter). CDK2-cyclin A complex at 10 mg mL^-1^ was mixed at a ratio of 1:1 with precipitant solution, 900 nl total drop size, using a mosquito robot (SPT Labtech). Plates were incubated at 4 °C for 2 days and then returned to 20 °C and grown for a further 5 days. FragLite soaking experiments of CDK2-cyclin A using the FragLite library was carried out in house. Crystals were transferred to a 96-well Imp@ct plate (Grenier Bio-One), pre-loaded with 25 mM compound prepared using 5 % DMSO in CDK2-cyclin A precipitant buffer (100 mM HEPES pH 7.0, 1.2 M (NH_4_)_2_SO_4_, 0.76 M KCl) and soaked for 2 hr. CDK2-cyclin A crystals were harvested in 6 M sodium formate solution and flash cooled in liquid nitrogen. Briefly crystals were transferred to cryovials, mounted on canes and stored in liquid nitrogen before transferring to UniPucks for shipment.

Crystallographic diffraction experiments were conducted at 100 K at the Diamond Light Source (DLS), Harwell Campus, Oxfordshire, UK. Images were automatically processed at DLS using AutoProc^61^ and Xia2 with Dials^62,63^. Programs from the CCP4i2 software suite^64^ accessed from the graphical user interface^65^ were used to import and process merged data. Initial phases and ligand dictionaries were generated for each dataset using isomorphous replacement with the incorporation of Phaser^66^ and AceDrg^67^ pipelines. Log-likelihood gain (LLG) anomalous signal maps were derived from molecular replacement using Phaser. A starting model of CDK2-cyclin A was obtained from PDB code 6GUC^60^, which adopts the human cyclin A2 numbering. Phases were further refined through iterative rounds of model building in COOT^68,69^ and refinement in REFMAC5^70^. Protein models were visualized using CCP4MG^71^ and UCSF Chimera^72^.

## Data and Software Availability

Crystal structures generated in this study were deposited in the Protein Data Bank (PDB) under the following accession codes:

| Structure | PDB Accession Code | Source |
| --- | --- | --- |
| Apo CDK2/Cyclin A | 6GUC | Wood *et al.* (2019) *Cell Chem. Biol.* |
| CDK-cyclin A-FL1 | 9ETP | This paper |
| CDK-cyclin A-FL2 | 9ETB | This paper |
| CDK-cyclin A-FL3 | 9ETA | This paper |
| CDK-cyclin A-FL4 | 9ET9 | This paper |
| CDK-cyclin A-FL5 | 9ET8 | This paper |
| CDK-cyclin A-FL6 | 9ET7 | This paper |
| CDK-cyclin A-FL7 | 9ET6 | This paper |
| CDK-cyclin A-FL8 | 9ET5 | This paper |
| CDK-cyclin A-FL9 | 9ET4 | This paper |
| CDK-cyclin A-FL10 | 9ET3 | This paper |
| CDK-cyclin A-FL11 | 9ET2 | This paper |
| CDK-cyclin A-FL12 | 9ET1 | This paper |
| CDK-cyclin A-FL13 | 9ET0 | This paper |
| CDK-cyclin A-FL16 | 9ESY | This paper |
| CDK-cyclin A-FL14 | 9ESZ | This paper |
| CDK-cyclin A-FL17 | 9ESX | This paper |
| CDK-cyclin A-FL18 | 9ESW | This paper |
| CDK-cyclin A-FL19 | 9ESV | This paper |
| CDK-cyclin A-FL20 | 9ESU | This paper |
| CDK-cyclin A-FL22 | 9ESS | This paper |
| CDK-cyclin A-FL23 | 9ESR | This paper |
| CDK-cyclin A-FL24 | 9ESQ | This paper |
| CDK-cyclin A-FL26 | 9ESP | This paper |
| CDK-cyclin A-FL27 | 9ESO | This paper |
| CDK-cyclin A-FL28 | 9ESN | This paper |
| CDK-cyclin A-FL29 | 9ESL | This paper |
| CDK-cyclin A-FL30 | 9ESK | This paper |
| CDK-cyclin A-FL31 | 9ESJ | This paper |

## Acknowledgements

This research was supported by the Medical Research Council (Grant References MR/N009738/1 and MR/V029142/1), and Cancer Research UK (grant reference C2115/A21421). IH was supported by a studentship from the Medical Research Council Discovery Medicine North Doctoral Training Partnership. NT position within Newcastle Drug Discovery is supported by Astex Pharmaceuticals.

We wish to thank i03 beamline staff at Diamond Light Source (Oxford) for excellent facilities and Dr Arnaud Baslé at Newcastle University for assistance with data management. For the purpose of open access, the authors have applied a Creative Commons Attribution (CC BY) licence to any Author Accepted Manuscript version arising from this submission.

## Author Contributions

**Ian Hope:** Conceptualization, methodology, investigation (protein production, FragLite screening and complex structure determination), writing original draft, review and editing.

**Natalie Tatum:** Writing original draft, figures, review and editing.

**Mathew Martin:** Conceptualization, investigation (FragLite design, and fragment screening by crystallography), review and editing.

**Michael Waring:** Conceptualization, investigation (FragLite design), review and editing.

**Jane Endicott:** Conceptualization, resources, supervision, and funding acquisition, review and editing.

**Martin Noble:** Conceptualization, resources, supervision, review and funding acquisition.

## Supplemental Information

Document S1: Supplementary Figures S1-S3, Supplementary Table S1,

Spreadsheet S1: Supplementary Table S2

## Supplementary Information

**Supplementary Figure S1.**
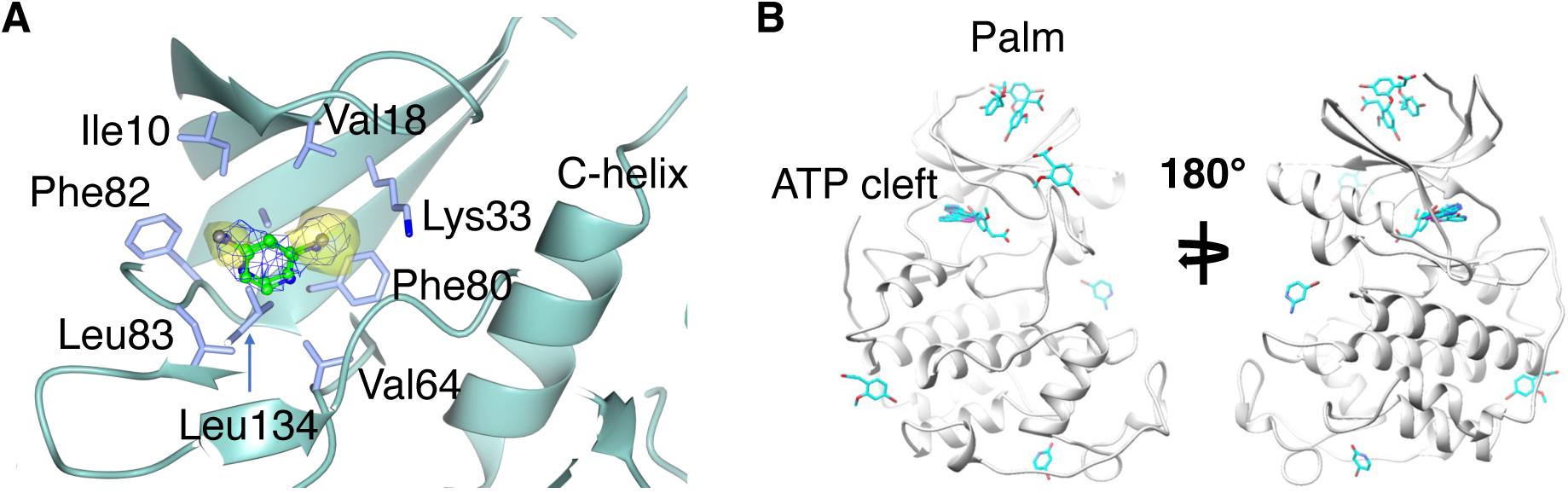
FragLite binding to CDK2-cyclin A and a comparison with the FragLite map of monomeric apo CDK2. **(A)** FL4 bound to the CDK2-cyclin A ATP-binding cleft. The Fo-Fc difference map (0.65 electrons/Å^3^) identifies the location of FL4 within the CDK2 ATP binding site, overlaid with the anomalous log-likelihood gain (LLG) map (anomalous signal shown in yellow) that identifies the bromine atom. Two alternative binding modes can be distinguished. **(B)** The FragLite set bound to monomeric CDK2 (Wood, Lopez-Fernandez, *et al.*, 2019).

**Supplementary Figure S2.**
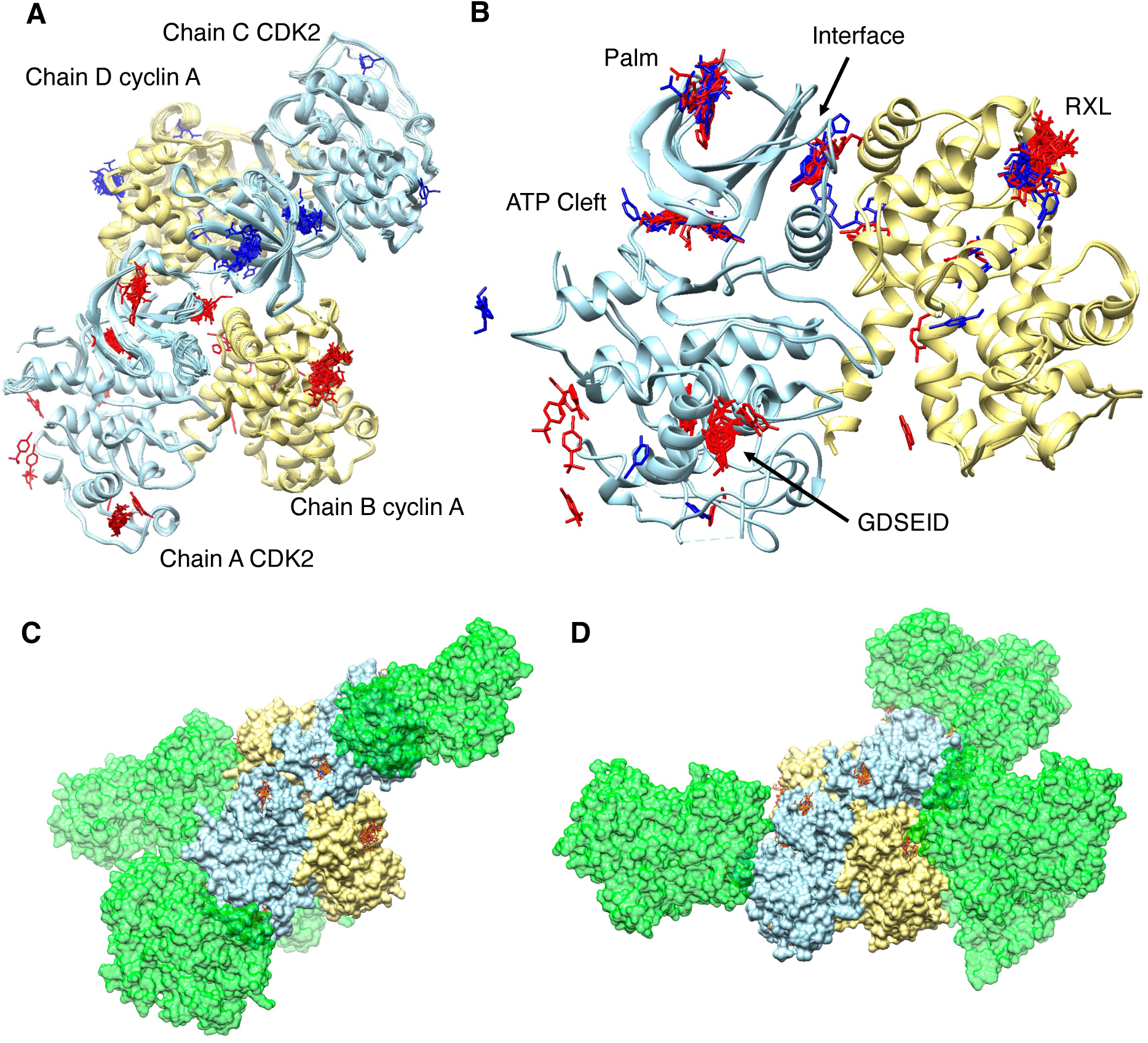
The dimer of dimers in the asymmetric unit allows for exposure of most of the protein surface. (**A**) The dimer of dimers for all 28 CDK2-cyclin A-FL structures superposed, with FragLites binding to the chain A/B monomer in red and FragLites binding to the chain C/D monomer in blue**. (B)** The consolidated map of FragLite binding demonstrates the palm, ATP, interface and RXL recruitment sites are occupied by FragLites from both dimers and fragment poses differ due to different crystal environments; however due to the crystal contacts which support lattice formation **(C)**, the GDSEID site is only populated by fragments from the A/B monomer. **(C)** Crystal-contacting dimers (within 4Å) are shown in green surface, for chains A/B and (**D**) C/D.

**Supplementary Figure S3.**
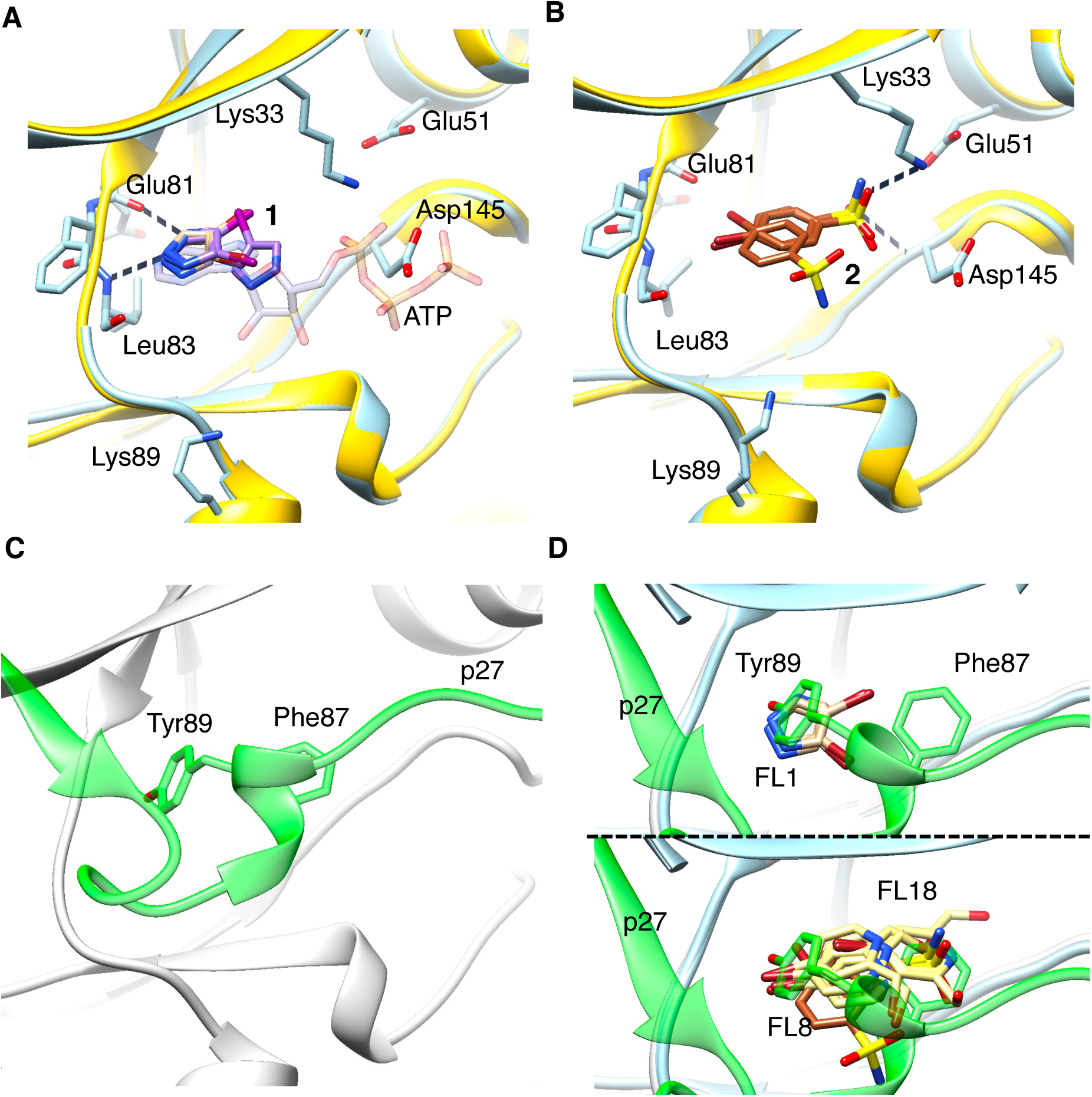
FragLite engagement at the ATP-binding cleft. **(A)** ATP-binding site mode 1. FL1 (carbon atoms in tan) and FL2 (carbon atoms in purple) occupy positions which overlay with the adenine moiety of ATP. The CDK2 fold extracted from CDK2-cyclin A bound to FL1 is drawn in blue ribbon. For comparison, the structure of CDK2-cyclin A bound to ATP (PDB 1FIN) is rendered in purple with transparent ATP. Hydrogen bonds are formed between FLs 1 and 2 with the backbone carbonyl of Glu81 (2.9 Å) and backbone amide of Leu 83 (2.8 Å). CDK2 residues 1-20 (the G-rich loop) have been omitted for clarity. **(B)** ATP site mode 2. FL8-bound structure shown in blue ribbon against ATP-bound CDK2 from PDB 1FIN in purple. FL8 (sienna carbons) exemplifies an alternative orientation which engages with polar residues Lys33 (hydrogen bond distance 2.7Å), Lys89, Glu51 and Asp145. CDK2 residues 1-20 (the G-rich loop) have been omitted for clarity. **(C)** Partner protein p27 (green ribbon) bound at the ATP cleft. Residues Phe87 and Tyr88 of p27 are the key interacting residues shown in stick representation. **(D)** Both FragLite binding modes mimic p27 engagement. For comparison, the structure of the CDK2-cyclin A-p27 complex (PDB 1JSU) is rendered in green ribbon (p27) and grey ribbon (CDK2). FL1 (upper panel) and FL8/18 (lower panel, FL8 in sienna carbons, FL18 in yellow carbons) occupy the same positions as p27 Phe87 and Tyr88 as shown in panel C.

**Supplementary Table S1.**
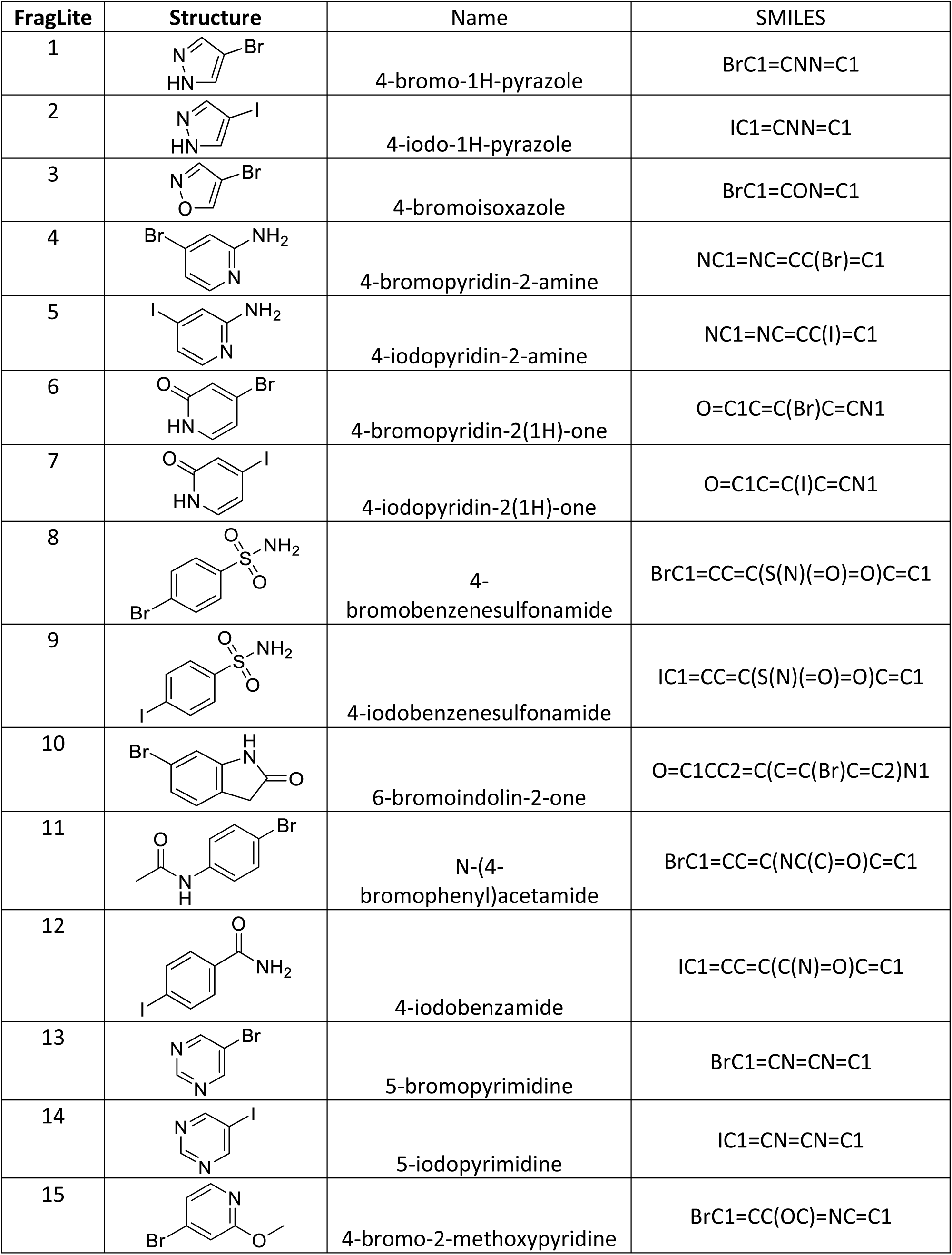
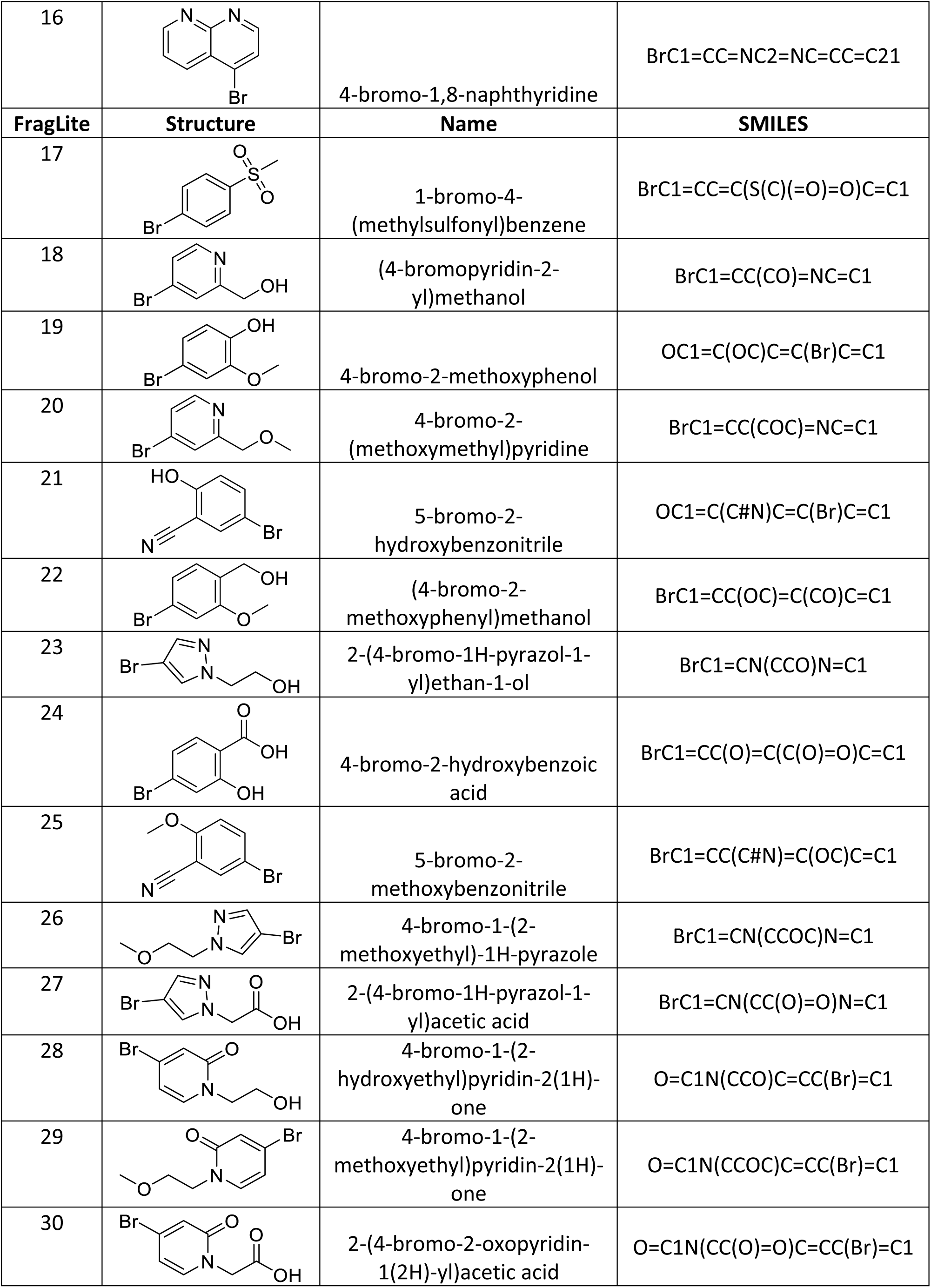
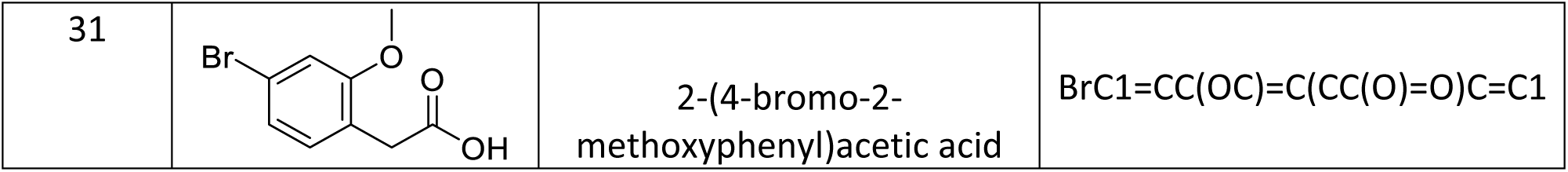

